# phyr: An R package for phylogenetic species-distribution modelling in ecological communities

**DOI:** 10.1101/2020.02.17.952317

**Authors:** Daijiang Li, Russell Dinnage, Lucas Nell, Matthew R. Helmus, Anthony Ives

**Affiliations:** Department of Wildlife Ecology & Conservation, University of Florida, Gainesville, FL 32611; Research School of Biology, Australian National University, Acton ACT 2601, Australia; Institute for Applied Ecology, University of Canberra, Bruce ACT 2617, Australia; Department of Integrative Biology, University of Wisconsin-Madison, Madison, WI 53706; Integrative Ecology Lab, Center for Biodiversity, Department of Biology, Temple University, Philadelphia, PA 19122

**Keywords:** phylogenetic diversity, phylogenetic generalized linear mixed models, functional trait, trait correlation, model-based methods, Joint Species Distribution Model

## Abstract

1. Model-based approaches are increasingly popular in ecological studies. A good example of this trend is the use of joint species distribution models to ask questions about ecological communities. However, most current applications of model-based methods do not include phylogenies despite the well-known importance of phylogenetic relationships in shaping species distributions and community composition. In part, this is due to lack of accessible tools allowing ecologists to fit phylogenetic species distribution models easily.
2. To fill this gap, the R package `phyr` (pronounced fire) implements a suite of metrics, comparative methods and mixed models that use phylogenies to understand and predict community composition and other ecological and evolutionary phenomena. The `phyr` workhorse functions are implemented in C++ making all calculations and model estimations fast.
3. `phyr` can fit a variety of models such as phylogenetic joint-species distribution models, spatiotemporal-phylogenetic autocorrelation models, and phylogenetic trait-based bipartite network models. `phyr` also estimates phylogenetically independent trait correlations with measurement error to test for adaptive syndromes and performs fast calculations of common alpha and beta phylogenetic diversity metrics. All `phyr` methods are united under Brownian motion or Ornstein-Uhlenbeck models of evolution and phylogenetic terms are modelled as phylogenetic covariance matrices.
4. The functions and model formula syntax we propose in `phyr` serves as a simple and unified framework that ignites the use of phylogenies to address a variety of ecological questions.

## Introduction

Ecological communities are collections of species that occur within the same geographical area. Which species occur within communities depends on the dispersal ability of species to enter the community, the environmental conditions that they find there, and the interactions that they have with other species in the community. These three processes – dispersal, environmental tolerance, and species interactions – depend on the traits that species possess and hence reflect evolutionary history and biogeographic processes (Warren, Cardillo, Rosauer, & Bolnick, 2014; Gerhold, Carlucci, Procheş, & Prinzing, 2018). For example, the larvae of an aquatic insect species might only occur in a lake if its adult stage has long-distance flight capabilities, if it can tolerate the low pH of the lake, and if it can avoid the predators that are common. Because traits play a central role in the composition of species that make up a community, community composition will likely reflect, at least in part, phylogenetic relationships among species. For example, two closely related insects might have similar dispersal capability, pH tolerance, and predator avoidance behavior, making them more likely to occur in the same lake. The recognition that phylogenetic relationships can increase our understanding of communities has led to a growing number of statistical methods for analyzing phylogenetic community composition (Losos, 1996; Webb, 2000; Webb, Ackerly, McPeek, & Donoghue, 2002; Cavender-Bares, Ackerly, Baum, & Bazzaz, 2004; Helmus, Savage, Diebel, Maxted, & Ives, 2007; Ives & Helmus, 2011; Frishkoff, Valpine, & M’Gonigle, 2017; Li et al., 2017).

Just as the distributions of two species might reflect their proximity on a phylogenetic tree, the species occurring at two sites might reflect the sites’ geographical proximity. The most immediate possible cause of spatial correlations in species distributions is dispersal, if nearby sites are more likely to be colonized by a species. Spatial proximity may also be a surrogate for environmental variables that are unknown or unmeasured. For example, an insect species might occur in two nearby lakes because they both have low pH, yet pH has not been measured. Just as phylogenetic relationships among species can generate correlations between species in which sites they occupy, so too can spatial proximity generate correlations between sites in the species they contain (Cressie, 1991; Ives & Zhu, 2006).

How species respond to environmental factors, and how they respond to each other, depend on their traits. Therefore, correlations among functional traits can provide insights about the evolutionary history that has shaped species traits so they can occupy the same sites. For example, two insect species that occur in the same lake might share both long-range flight abilities and tolerance to low pH. Is the positive correlation between these two traits caused by correlated selective forces? A challenge to answering this question is that phylogenetic correlations between trait values might reflect species phylogenetic relatedness rather than shared selection: two species might have both long-range flight abilities and tolerance to low pH only because they are phylogenetically closely related. To distinguish between these two explanations – convergence of suites of traits due to shared selective forces versus similarity due to phylogenetic relatedness – it is necessary to account for phylogenies when performing correlation analyses between traits that could explain similarities in the distributions of species.

Statistical models for phylogenetic community composition provide flexible tools for exploring the many possible factors underlying the distribution of species and the composition of communities (Ives & Helmus, 2011; Ovaskainen & Soininen, 2011; Warton et al., 2015). The models can describe complex relationships in the data, such as how phylogenetically related species might respond similarly to the same environmental gradient, or how phylogenetically related species might exclude each other from the same communities. They also give a firm statistical basis to test these patterns, the ability to simulate data sets from the fitted model, and the ability to predict the composition of unsurveyed communities. These benefits of phylogenetic community composition models come with costs: building models can be intricate and fitting them computationally slow.

The R package `phyr` is designed to overcome many of these costs with a user-friendly interface, flexibility to build a rich collection of models, and good computational performance. Below, we first give a brief overview of the structure and syntax of two key functions `pglmm()` and `cor_phylo()`. `pglmm()`allows the formulation of a diverse set of phylogenetic generalized linear models (PGLMM) that can be used not only to analyze phylogenetic community composition but also comparative models for Gaussian and non-Gaussian data. `cor_phylo()` computes the Pearson correlations among species traits while simultaneously estimating the strength of phylogenetic signal within each trait. We then compare `pglmm()` and `cor_phylo()` to methods and programs that are currently available. Finally, we apply `pglmm()` and `cor_phylo()` to simulated data to illustrate their implementation and output.

## Overview of phyr

`Phyr` contains three groups of functions (Table 1): phylogenetic generalized linear mixed models (`pglmm()`), phylogenetic comparative methods (`cor_phylo()` and `pglmm_compare()`), and community phylogenetic diversity metrics (e.g., `psv()`, `pse()`). The workhorse functions of all groups are written in C++ to increase computational speed. Here, we will focus on the first two groups of functions (especially `pglmm()` and `cor_phylo()`), because they are more complicated and less readily available to practitioners than community phylogenetic diversity metrics.

**Table 1:** List of main functions in the phyr package.

| Group | Main Functions | Brief Description |
| --- | --- | --- |
| Mixed Models | `pglmm()` | Phylogenetic Generalised Linear Mixed Model for ecological community data (e.g., species composition across sites; bipartite interactions) |
| Comparative Methods | `cor_phylo()` | Correlations among multiple traits with phylogenetic signal |
|  | `pglmm_compare()` | `pglmm()` tailored for comparative data in which species (tips of a phylogeny) only occur once |
| Metrics | `psv()`; `pse()`; `psr()`; `psc()`; `psd()` | Phylogenetic alpha diversity of communities |
|  | `pcd()` | Pairwise phylogenetic beta diversity of communities |
|  | `v cv2()` | Convert a phylogeny to a covariance matrix, a faster version of `ape::vcv()` |

### pglmm()

Function `pglmm()` constructs and fits generalized linear mixed models that incorporate covariance matrices containing the phylogenetic relationships among species. The syntax for `pglmm()` resembles that used in the R package `lme4` (Bates, Mächler, Bolker, & Walker, 2015), and indeed `pglmm()` will fit most of the models that can be fit with `lmer()` and `glmer()`. `pglmm()` goes beyond `lmer()`and `glmer()` by allowing the specification of covariance matrices, which could be phylogenetic covariance matrices or any other covariance matrices that the user defines (e.g., spatial or temporal autocorrelation matrix). `pglmm()` can also fit models with “nested” covariance structures (e.g., a species phylogenetic covariance matrix nested within a site covariance matrix). `pglmm()` can operate in both frequentist mode, with the distribution of species among communities being Gaussian, binary, binomial or Poisson, and Bayesian mode with the addition of zero-inflated binomial and Poisson distributions. Finally, it is our hope that the formula syntax of `pglmm()` can be used to fit similar models with other programs such as Stan (e.g. via R package `brms` Bürkner, 2018).

A general example of the syntax for `pglmm()` is

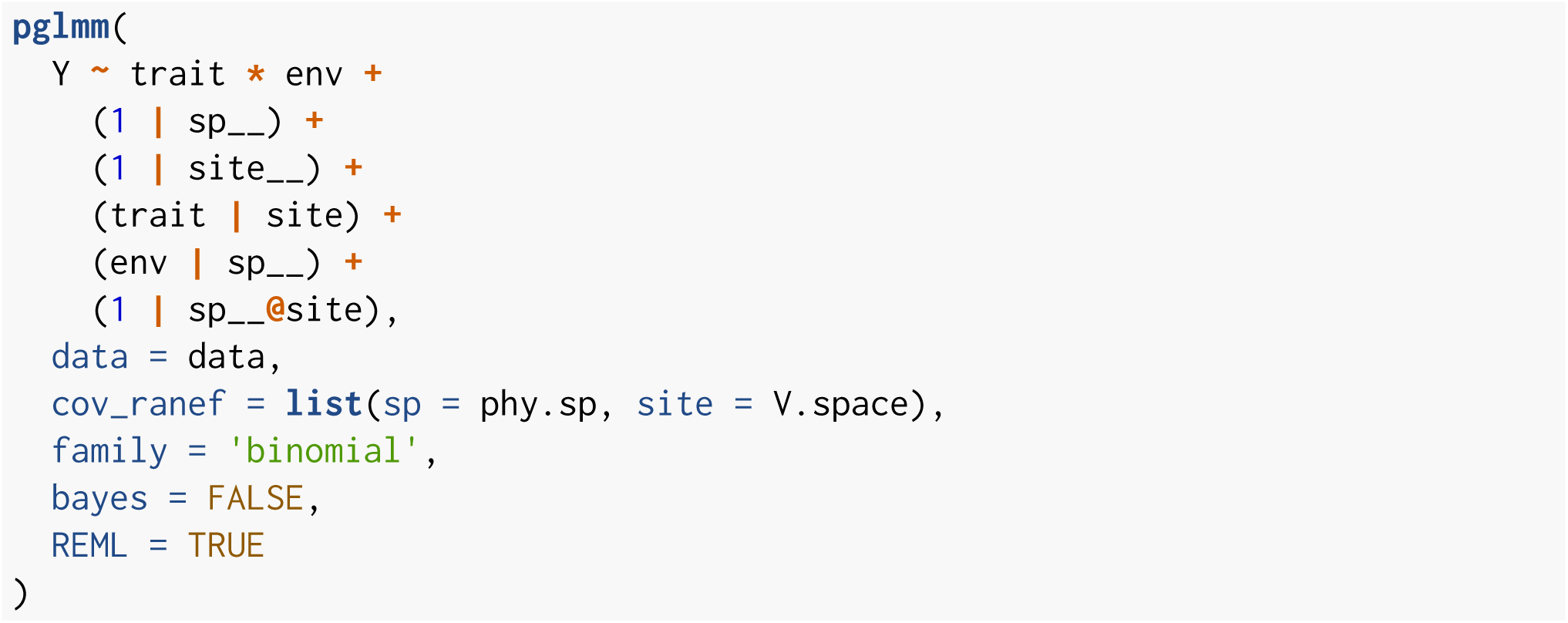

Here, Y is a binary (Bernoulli) dependent variable which takes values of either 0 or 1. The specification `family = ‘binomial’` allows binary data and also binomial data for which Y is a matrix containing columns for successes and failures. The independent variables `trait` and `env` take on different values for each species and site, respectively. Sites (`site`) and species (`sp`) are treated as random effects: `(1|site)` implies that a value from a Gaussian random variable is picked for each site, thereby representing unmeasured differences among sites. For the case of species, the double underscore in `(1|sp__)` implies that, in addition to a random effect for species, there is a second random effect which contains the phylogenetic relationships among species (or some other correlation structure specified by the user). The phylogenetic random effect assumes that values for each species are picked from a multivariate Gaussian distribution with phylogenetic covariance matrix Σ. A covariance matrix Σ is specified by `cov_ranef = list(sp = phy.sp, site = V.space)`. The covariance matrix `phy.sp` associated with species can be a `phylo` object from the R package `ape` (Paradis & Schliep, 2018). To construct from a ‘phylo’ object, `pglmm()` assumes that the residual variation associated with species follows a Brownian motion model of evolution, so that the covariance between species is proportional to their shared evolutionary history (e.g., shared branch length on a phylogeny). It is also possible to specify an explicit covariance matrix, such as `site = V.space`, where `V.space` is a covariance matrix created from the distance between sites.

The syntax `(1|sp__)` or `(1|site__)` generates two random effects, one without and one with phylogenetic or spatial covariances; in contrast, `(1|sp)` would generate only a single random effect that is independent among species. `pglmm()` forces in a term for `(1|sp)` whenever `(1|sp__)` is specified, because otherwise any difference among species would be captured by the diagonal elements in Σ even in the absence of covariances among phylogenetically related species which are specified by the off-diagonal elements of Σ. Therefore, if `(1|sp)` were not included, this could lead to the identification of phylogenetic signal in the abundances of species even in its absence from a community. To account for differences among sites in how they select for species with different traits, `(trait|site)` allows the slope of Y against `trait` to be a Gaussian random variable. Similarly, to account for the differences among species for how they respond to `env`, `(env|sp__)` allows the relationship of Y against `env` to be given by two slopes, the first slope that is picked from a Gaussian random variable in which species are independent and the second slope that is picked from a multivariate Gaussian with covariance matrix Σ. Finally, (1|sp__@site) generates a nested term: within a site, the residual variation in Y shows phylogenetic relatedness, with phylogenetically related species more likely to occur in the same site. Note that `(1|sp__)` differs from (1|sp__@site) because `(1|sp__)` generates differences in the mean value of Y for species across all sites, whereas (1|sp__@site) is local to sites, giving the covariances among species only within sites. This nested term can be used to test for community clustering or overdispersion (Webb et al., 2002; Ives & Helmus, 2011). Other forms of a nested term are available in `pglmm()`, which can be used to study more complicated questions such as bipartite networks.

With `bayes = FALSE`, `pglmm()` is fitted using a frequentist approach. ML or REML is used for fitting, with `REML = TRUE` as the default. For a non-Gaussian model (e.g., `family = ‘binomial’`), an iterated quasi-likelihood method is used for model fitting which gives the approximate likelihood; p-values for the fixed effects are given by a Wald test and for the random effects by profile likelihood, although we recommend bootstrap-based tests when computationally feasible. Note that `REML = TRUE` is an option for non-Gaussian models (in contrast to `glmer()`) due to the algorithm used. With `bayes = TRUE`, a Bayesian approach is implemented using INLA (Rue, Martino, & Chopin, 2009), which gives parameter estimates and credible intervals. For large problems with the number of species-site combinations exceeding 2000, the Bayesian computations are considerably faster than the frequentist computations. Finally, a key to interpreting the results from a model is understanding the structure of the covariance matrices associated with the random effects. Therefore, `pglmm()` has associated plotting functions `pglmm_plot_ranef()` that present the design matrices for the random effects (Fig. 1).

**Figure 1:**
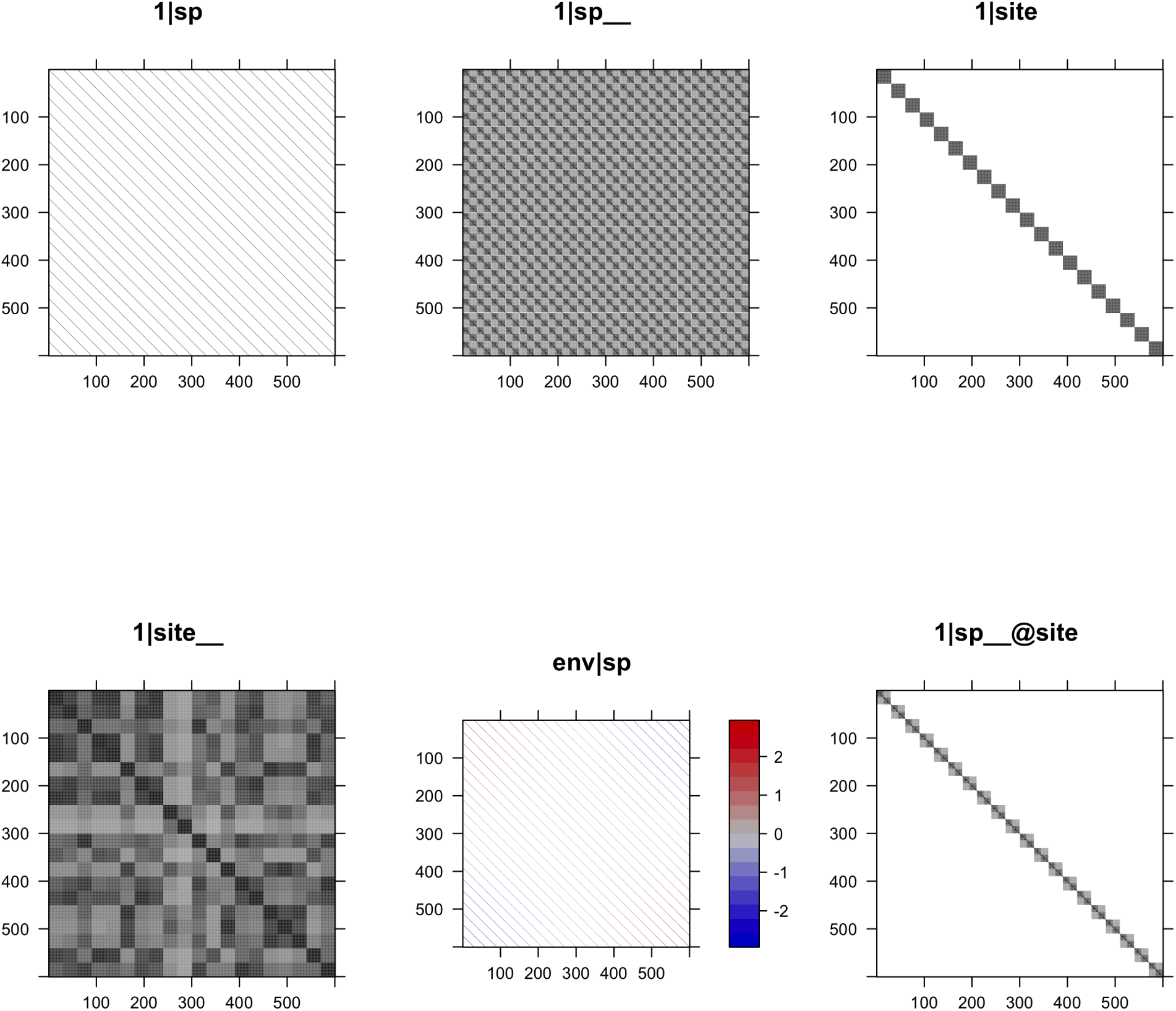
The structures of design matrices of random terms in a phylogenetic generalized linear mixed model with 30 species and 20 sites. Random terms `1|sp` and `1|site` allow different species or sites to have different intercepts, respectively. Random terms `1|sp__` and `1|site` allow closely related species or sites with similar conditions to have similar intercepts, respectively. Random term `env|sp` allows different species to have different environment-abundance relationships independently. Random term 1|sp__@site is a nested term and allows closely related species more likely to occur in the same site.

Whereas `pglmm()` is designed to accept community composition data, in which the same species can occur in multiple sites, the algorithm used by `pglmm()` can equally be used for comparative data in which each species is represented by only a single data point. `pglmm_compare()` is a wrapper for `pglmm()` that is tailored for comparative data and thus provides an easy-to-use function for analyzing non-Gaussian phylogenetic data.

#### cor_phylo()

`cor_phylo()` makes it possible to compare suites of traits among species, accounting for their phylogenetic relatedness (Zheng et al., 2009; Johnson, Ives, Ahern, & Salminen, 2014). To identify suites of traits under joint selection, such as traits that together make up adaptive syndromes, it is necessary to perform a correlation analysis in which phylogenetic relatedness is factored out. `cor_phylo()` does this. It can also include within-species variation (e.g., measurement error) which should better-expose the underlying correlations in traits among species. Whereas `pglmm()` can be used to identify the composition of communities within a region, `cor_phylo()` can be used to assess patterns of traits among species that make up the regional species pool.

The syntax for `cor_phylo()` is

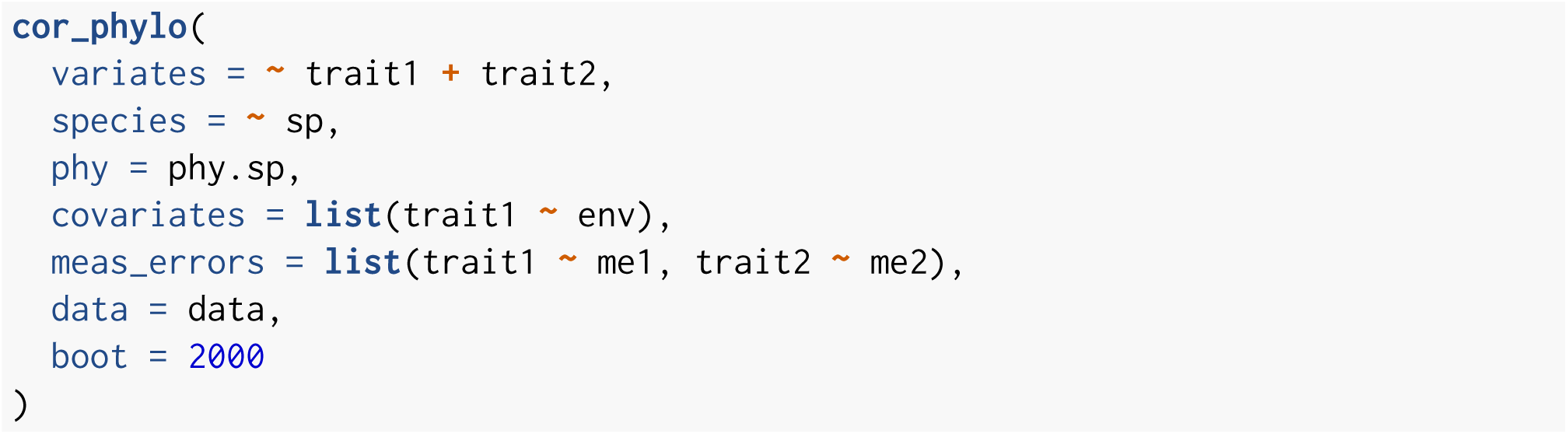

In this example, the correlation between `trait1` and `trait2` is computed, and the column named `sp` in `data` identifies the species. The object `phy.sp` specifies the phylogenetic covariance matrix as a ‘phylo’ object from the `ape` package. `cor_phylo()` estimates the phylogenetic signal for each trait by assuming that trait evolution is given by a Ornstein-Uhlenbeck process. The term `covariates = list(trait1 ∼ env)` includes the independent variable `env` for `trait1`, to remove possible confounding effects; only an intercept is estimated if no covariate is provided for a trait. Within-species variation is specified by `meas_errors = list(trait1 ∼ me1, trait2 ∼ me2)`, where `me1` and `me2` are the standard errors for `trait1` and `trait2`, respectively, of values at the tips of the phylogenetic tree. If within-species standard errors are not provided for a given trait, the trait values are assumed to be known without error. Finally, `cor_phylo()` can perform parametric bootstrapping to give confidence intervals for all parameter estimates: correlations, phylogenetic signals, covariate coefficients, and coefficient covariances.

## Relationships to other methods and software

Earlier versions of `pglmm()` and `cor_phylo()` both appear in existing R packages (`pez` (Pearse et al., 2015) and `ape` (Paradis & Schliep, 2018), respectively), although the versions in `phyr` represent considerable improvements in ease-of-use, computational speed, and flexibility. Both have new syntax that makes them more intuitive to use. `pglmm()` also has new associated functions that plot the design of the covariance matrices (Fig. 1), making model interpretation easier. Both are now coded in C++ (for key functions), which speeds computation time by 5-10X. `pglmm()` now supports several non-Gaussian distributions and allows Bayesian analyses using INLA (Rue et al., 2009) that is particularly useful for large datasets. Finally, both include more output; for example, both now include facilities to perform likelihood ratio tests and compute AIC and BIC values for model comparisons.

### pglmm()

`pglmm()` is syntactically modeled after `lmer()` and `glmer()` in `lme4` (Bates et al., 2015), although it allows the specification of phylogenetic covariance matrices. `pglmm()` also allows “nested” models (with block-diagonal covariance matrices) which arise when phylogenetic covariances only act within single communities, rather than among communities; an example is illustrated by the (1|sp__@site) term in Fig. 1. Such nested models make it possible to assess whether phylogenetic relatedness affects the abundance of species within the same communities, such as whether competition between closely related species excludes one of the competitors from communities where the other is present. Nested models are structurally incompatible with the architecture of `lme4`.

There are alternative programs to `pglmm()`, although they have limitations that `pglmm()` overcomes. Hadfield, Krasnov, Poulin, & Nakagawa (2013) use the R package `MCMCglmm` (Hadfield, 2010) to perform phylogenetic community analyses, although they also use ASReml because its penalized quasi-likelihood (PQL) approach is computationally much faster. Hierarchical Modelling of Species Communities (HMSC-R) (Tikhonov et al., 2019) performs community analyses using Bayesian MCMC approaches, although it does not include nested terms. It is also possible to code specific phylogenetic community models using flexible Bayesian platforms such as WinBugs, Stan, and JAGS, although this will involve considerable programming and expertise.

#### PGLMM as a Joint Species Distribution Model (JSDM)

Joint Species Distribution Models (JSDMs) are models where the response variable is distribution (abundances or occurrences) of multiple species across sites or samples, where all species are modeled jointly, usually by allowing non-zero covariance between either species-level errors, species-level coefficients in the model, or both (Warton et al., 2015). `pglmm()` in `phyr` is a joint species distribution model where the (residual) dependencies among species are modeled in a way that incorporates phylogenetic relatedness. JSDMs, and Species Distribution Models (SDM) in general, have typically been focused on producing accurate predictions of how species are distributed, usually in a geographic context. However, they can also be used for making inferences about the biology of species, such as which environmental factors are important in explaining the distribution of a species or set of species, and whether traits or evolutionary history can help explain these distributions. It is this kind of inference that is the focus of the JSDM implemented in `pglmm()`. There is often a trade-off between improving predictions and making solid inferences, because increasing the complexity or flexibility of a model can improve its predictive power, but this same complexity makes it more difficult to understand what biology is being represented by the model outputs. By incorporating phylogenetic information, `pglmm()` has two uses. First, by identifying correlations that might be expected among species due to phylogeny, `pglmm()` gives better statistical properties for tests of factors underlying community composition. For example, Li & Ives (2017) show that failure to account for phylogenetic correlations can inflate type I errors in tests for associating environmental variables and traits that underlie community composition. Second, `pglmm()` allows explicit focus on the importance of evolutionary history in structuring species assemblages, since phylogenetic covariances are explicitly estimated. This is in contrast to many other JSDMs (e.g. those in described in Wilkinson, Golding, Guillera-Arroita, Tingley, & McCarthy, 2019), which attempt to estimate all pairwise species covariances without accounting for phylogeny.

Of course, the goals of solid inference and prediction are not mutually exclusive. Good prediction requires avoiding overfitting, which can be facilitated by reducing the number of parameters in the model. In some systems, it may be possible to make better predictions using a simple phylogenetic model if phylogeny is a strong predictor of species covariance, or if many species are poorly sampled and thus estimating covariances between them individually results in higher prediction variance. Using phylogeny can help closely related species share statistical strength through phylogenetic partial pooling. Ultimately, it can be powerful to use the same kind of statistical framework to do both predictive and inferential work in ecology, because it allows ecologists to smoothly move between these two goals and more easily and quickly draw mutual insights between them.

### cor_phylo()

The R package `mvMORPH` (Clavel, Escarguel, & Merceron, 2015) can fit a broad range of models, of which `cor_phylo()` can be formulated as a special case. While `cor_phylo()` does not have the flexibility of `mvMORPH`, it is correspondingly simpler to use. Also, `cor_phylo()` has built-in bootstrapping capabilities that are necessary to give confidence in the parameter estimates and p-values. The function `evolvcv.lite()` in the R package `phytools` (Revell, 2012) will compute phylogenetic correlations, and changes in phylogenetic correlations through time (see also Caetano & Harmon, 2018), although the phylogenetic covariance matrix is derived under the assumption of Brownian motion evolution. This contrasts `cor_phylo()` in which the strength of phylogenetic signal is computed at the same time as the correlation. It is also possible to code the `cor_phylo()` model using platforms such as WinBugs, Stan, and JAGS; but again, this will require considerable programming and expertise.

## Example usage

We simulated datasets to demonstrate how to use `pglmm()` and `cor_phylo()`. Details about simulations of PGLMM can be found in the Appendix. Our goal in this section is to provide some general ideas about the inputs and outputs of these two functions instead of testing their statistical performances or interpreting the ecological meanings of model results. For those purposes, please see the package vignettes and Ives (2018).

### pglmm()

We fitted a PGLMM that examined how a hypothetical functional trait, environmental gradient, and their interaction affect distributions of 30 species across 20 sites. We focused on abundance and used the default family of data distribution (Gaussian), but other distributions can also be specified by resetting the `family` argument. Phylogenetic relationships among species and site spatial autocorrelations are specified by `cov_ranef = list(sp = phy, site = V.space)` where `sp` and `site` are group variables of random terms, `phy` can be a phylogeny with class `phylo` or a phylogenetic covariance matrix, `Vspace` is a covriance matrix among sites. This model can also be fitted with Bayesian framework by setting `bayes = TRUE`, which is recommended when data set is large.

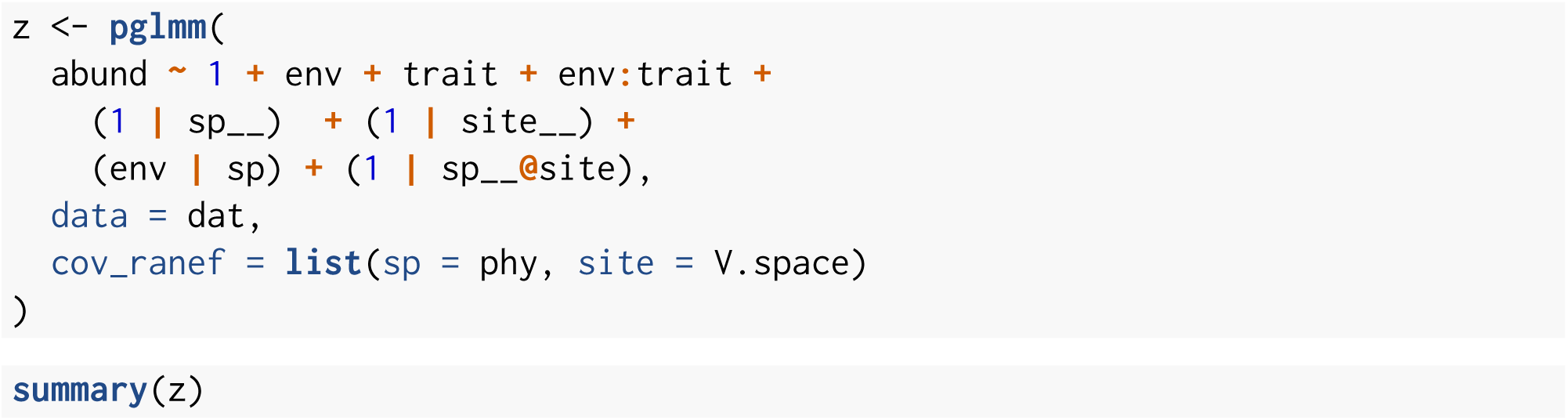

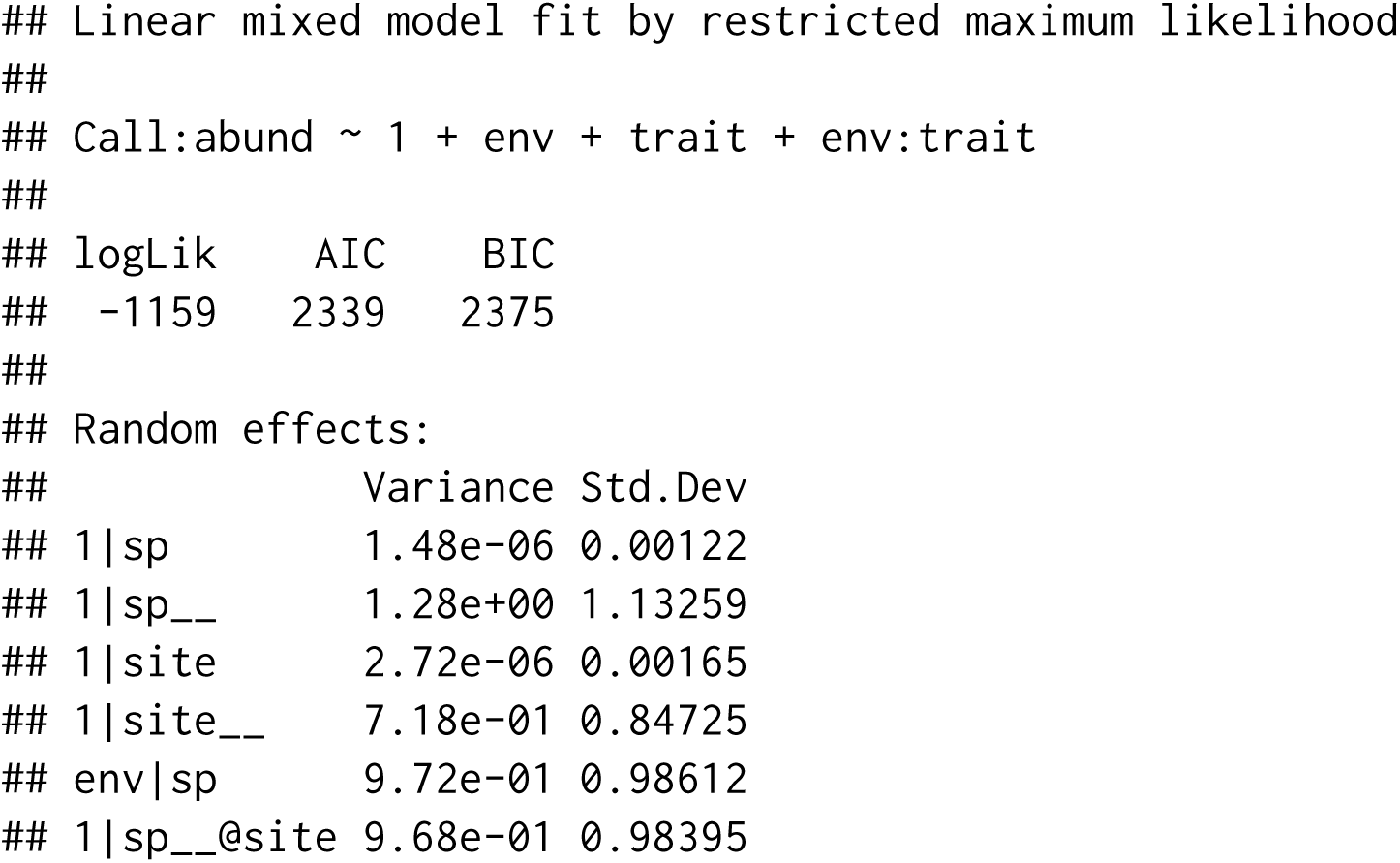

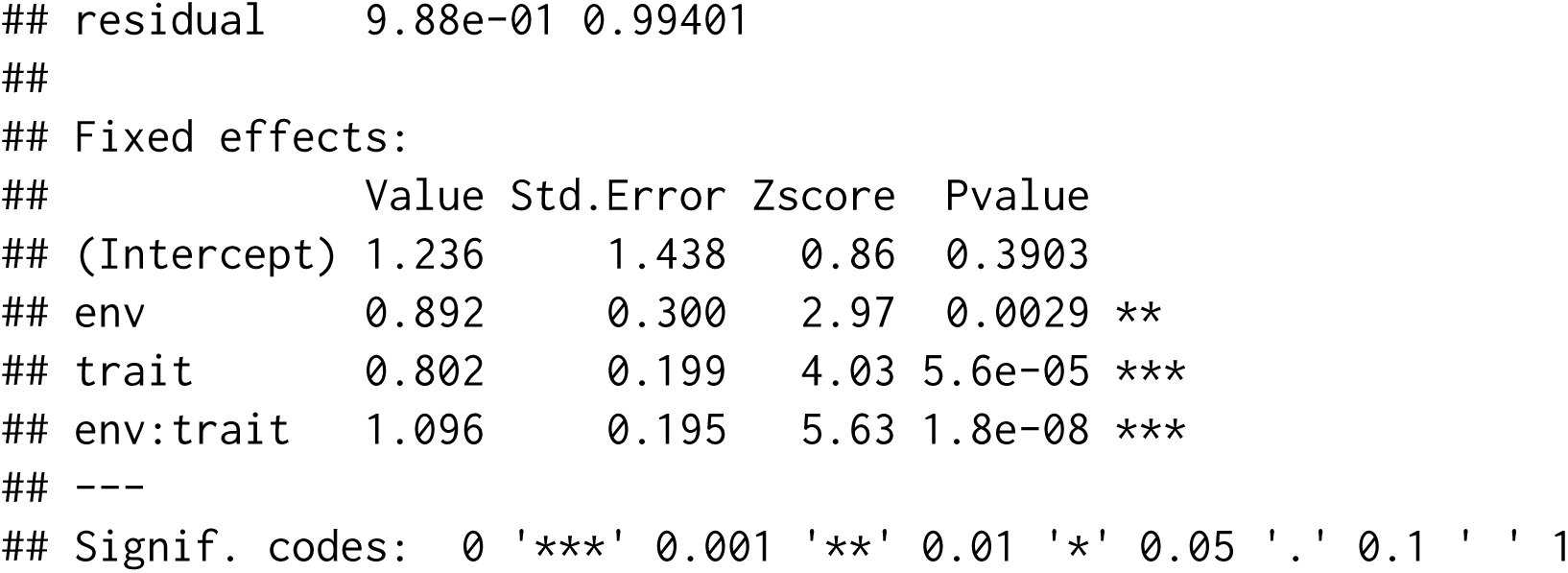

The summary of model results includes the model fitting method (maximum likelihood or bayesian), the model formula, log likelihood and other related statistics (AIC, BIC, DIC), estimates of variances of random terms, coefficients of fixed terms and their uncertainties. These results show that `pglmm()` correctly recovered that the hypothetic functional trait interacted with environmental variable to affect species composition.

### cor_phylo()

Here, we simulated two hypothetical functional traits (`trait_1` and `trait_2`) for 50 species. We set the true correlation between these two traits to be 0.7 and their phylogenetic signals to be 0.3 and 0.95, respectively. We also set their measurement errors to be 0.2 and 1, respectively.

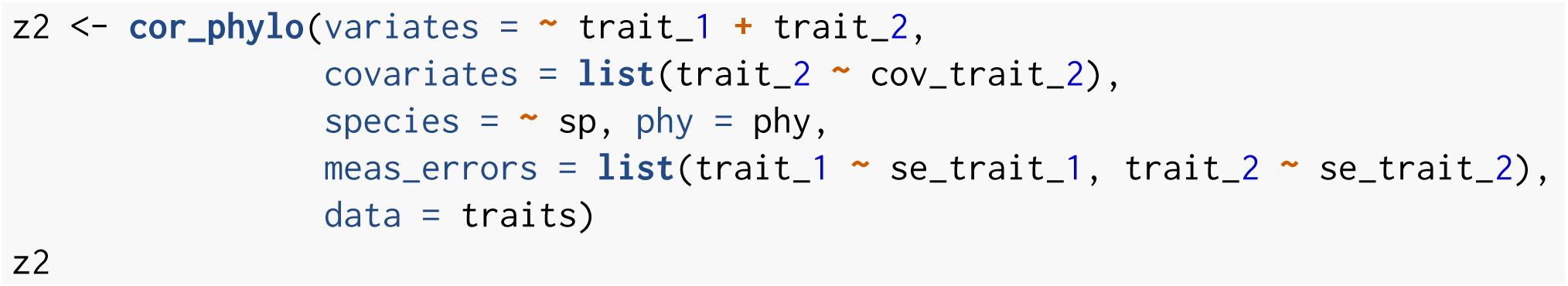

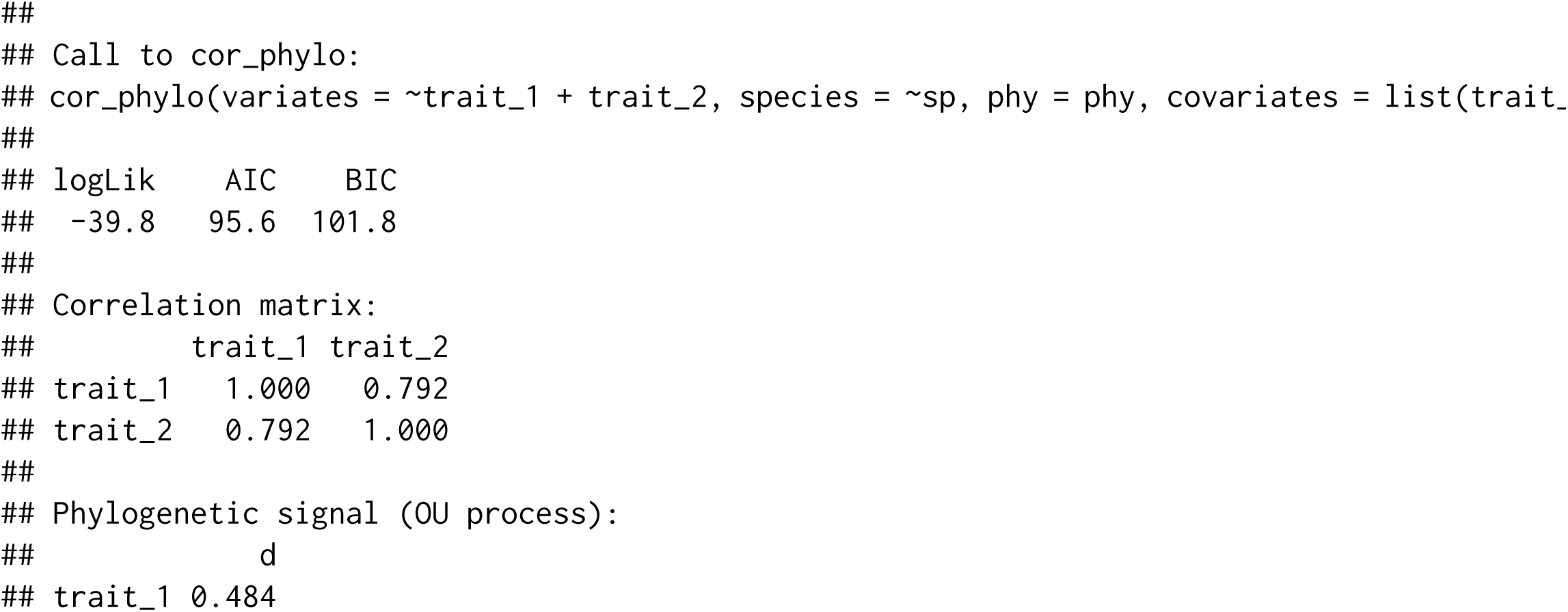

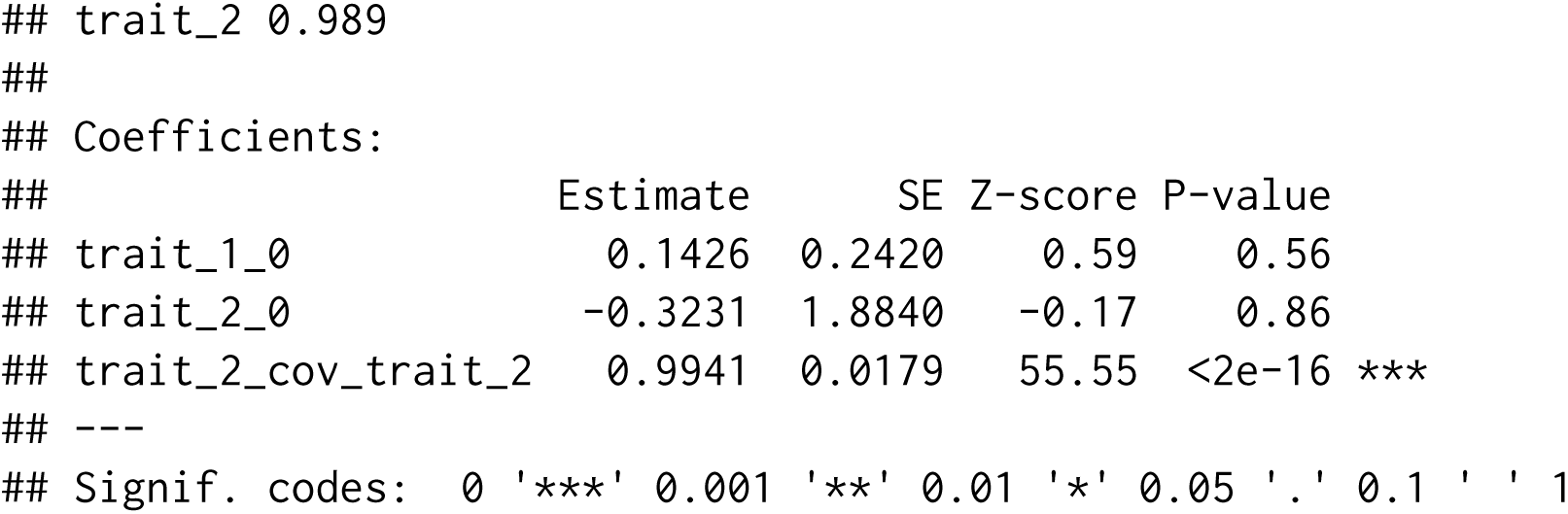

The output of `cor_phylo()` includes log-likelihood values, AIC, BIC, estimated correlation matrix of traits, estimated phylogenetic signals of traits, estimated coefficients and their uncertainties (standard errors, Z scores, and p values). In this example, the model gave good estimates of the parameters used to simulate the data. If bootstrapping was enabled by setting the `boot` argument, the lower and upper boundaries of correlations, phylogenetic signal values, and coefficients will be appended.

## Closing remarks

In recent years, there has been increasing effort to apply model-based approaches in community ecology. Despite the well-known importance of phylogenetic relationships in structuring species distributions and community composition, relatively few studies have incorporated phylogenetic relationships in model-based analyses of species distributions and community ecology. A potential reason is the lack of easy-to-use tools to facilitate the use of phylogenetic species-distribution modeling in ecological communities. The package `phyr` fills this gap by providing implementations of phylogenetic species-distribution models with flexible model formula syntax (`pglmm()`). It also includes other model-based functions that are useful for ecological studies such as estimating correlations among functional traits while accounting for their evolutionary history (`cor_phylo()`) and calculating community phylogenetic diversity (e.g. `psv()`) (Table 1).

The model formula of `pglmm()` is general and can be applied using other tools to fit phylogenetic species-distribution models. Thus, `pglmm()` can serve the developer community as a shell for new methods that fit GLMMs, with `phyr` providing an easy user interface. Using INLA as a backend to fit a Bayesian version of the PGLMM model is an example of this approach. To facilitate this end, we are developing phyr openly on github and actively encourage community contribution. We hope that the `phyr` package will help current and future researchers formulate and analyze phylogenetic species-distribution models.

## Acknowledgements

Funding for this work was provided by the National Science Foundation (US-NSF-DEB Dimensions of Biodiversity, 1240804).

## Authors’ contributions

D.L and A.R.I conceived the idea. All authors wrote the software and package documentations. All authors wrote the manuscript.

## Data Accessibility

No data were used in this study. R code used to simulate data for PGLMMs as described in the Appendix is available at https://github.com/daijiang/phyr_ms/blob/master/simulation_pglmm.R. R code used to simulate data for comparative methods is available at https://github.com/daijiang/phyr_ms/blob/master/simulation_cor_phylo.R. `phyr` is available at Github (https://github.com/daijiang/phyr) and CRAN (https://cran.r-project.org/package=phyr).

## Appendix

To demonstrate the usage of main functions in `phyr`, we simulated a dataset with 30 species and 20 communities. We first simulated a coalescent phylogeny of 30 species with function `ape::rcoal()`. For each species, we then simulated one continuous functional trait along the phylogeny. We also simulated one environmental variable with all 20 communities located evenly along the gradient. The environmental variable, functional trait, and their interaction all determine the abundance of species among sites according to the model below:

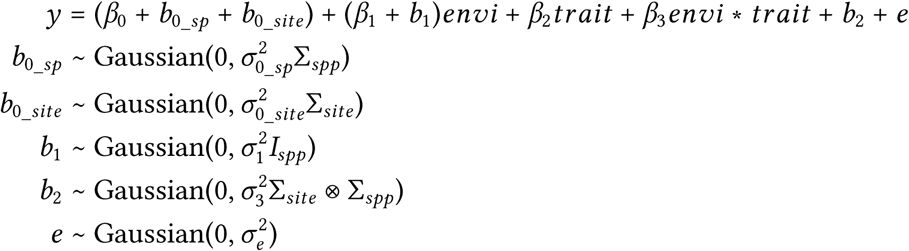

We set all coefficients (*β*_0_ to *β*_3_) to 1; we also set variances of all random terms to 1. Σ_*spp*_ is a covariance matrix converted from the phylogeny. Σ_*site*_ is a covariance matrix among sites, which was converted from the site spatial distance matrix. *I*_*spp*_ is the identity matrix so that we treat species as independent replicates. Σ_*site*_ ⊗ Σ_*spp*_ is a matrix generated as a kronecker product, and it makes closely related species more likely to be observed in the same site (i.e. underdispersion). Different species have different overall abundance (intercept) but closely related species (*b*_0_*sp*_) and similar communities (*b*_0_ *site*_) have similar overall abundance. Abundances of different species also change differently along the environmental gradient.

For demonstration purposes, we only simulated one functional trait and one environmental variable; in real datasets, multiple environmental variables and multiple functional traits can be included in the model.

